# Spectrum: Fast density-aware spectral clustering for single and multi-omic data

**DOI:** 10.1101/636639

**Authors:** Christopher R. John, David Watson, Michael Barnes, Costantino Pitzalis, Myles J. Lewis

## Abstract

Clustering of single or multi-omic data is key to developing personalised medicine and identifying new cell types. We present Spectrum, a fast spectral clustering method for single and multi-omic expression data. Spectrum is flexible and performs well on single-cell RNA-seq data. The method uses a new density-aware kernel that adapts to data scale and density. It uses a tensor product graph data integration and diffusion technique to reveal underlying structures and reduce noise. We developed a powerful method of eigenvector analysis to determine the number of clusters. Benchmarking Spectrum on 21 datasets demonstrated improvements in runtime and performance relative to other state-of-the-art methods.

## Introduction

Precision medicine is the concept that patients may be stratified into different classes to personalise therapy. A growing number of studies stratify patients using their genome wide expression data (e.g. mRNA, miRNA, protein, methylation), such as those by The Cancer Genome Atlas (TCGA)^1–7^ and other large consortia^8^. Clustering algorithms are used to find patient classes and may be run on data from single or multiple platforms. Single-omic (not including single cell RNA-seq) cluster analysis is performed by algorithms such as: Monte Carlo consensus clustering (M3C)^9^, CLEST^10^, PINSPlus^11^, and Similarity Network Fusion (SNF)^12^. However, clustering multi-omic data into an integrated solution is a major current challenge, state-of-the-art methods include: iClusterPlus^13^, SNF, CIMLR^14^, and PINSplus. The aims of multi-omic clustering are identifying a shared structure between platforms and reducing noise from individual platforms. These are two areas where there is scope for improvement on existing methods. Also important is improving runtimes and developing methods that are effective at both single and multi-omic cluster analysis.

Single-cell RNA-seq is a technique that can be used to detect new cell types by clustering of individual cell transcriptomes^15^. Analysing transcriptomes of individual cells has potential for furthering our understanding of biology and clinical applications. To date, single-cell RNA-seq clustering tools have been developed for clustering this data type only, however, this clustering problem is simply another type of single-omic data clustering. Differences with single-cell RNA-seq data include there are more samples and there are often many dense globular clusters. Tools applied in this domain include: single cell consensus clustering (SC3)^15^, Seurat^16^, MUDAN, and single-cell interpretation via multikernel learning (SIMLR)^17^. Maintaining fast runtimes is particularly important given the high samples sizes. SIMLR uses a sophisticated procedure to learn the optimal similarity matrix. However, it is quite time consuming and it is not clear if SIMLR provides clustering performance advantages relative to other methods. Given the importance of the above-mentioned clustering tasks, it is timely to develop an effective solution that can address any of these challenges.

Spectral clustering refers to a large class of algorithms that have grown rapidly in the machine learning field due to their ability to handle complex data^18–22^. They are characterised by clustering eigenvectors derived from a matrix representing the data’s graph^18^. A few of these methods are currently applied in genomic data analysis^12,17^. However, there have been several advancements in spectral clustering that provide opportunities for new method development. A key development has been the density-aware kernel^22^, which calculates sample density to strengthen connections in high density regions of the graph. A recent integrative method uses tensor product graph (TPG) integration and diffusion to leverage higher order information beyond that provided by each individual data source and reduce noise^19^. Another method retrieves eigenvectors of the data’s graph selected according to their multimodality for Gaussian Mixture Modelling (GMM) with the Bayesian Information Criterion (BIC) to decide on the number of clusters (K)^20^. The Fast Approximate Spectral Clustering (FASP) method^23^ enables rapid analysis of hundreds of thousands of samples on a desktop computer. Inspired by this work our aim was to assemble and advance it by developing Spectrum.

Spectrum includes both methodological advancements and implements pre-existing techniques. Highlights of Spectrum, that make it distinct from previous solutions^12,17^, include: 1) A new self-tuning kernel that is density- and scale-aware; 2) A TPG data integration and diffusion procedure to combine data sources; 3) Implementation of the FASP method for massive datasets; 4) A new technique based on eigenvector distributional analysis to find the optimal K; 5) A nonlinear dimensionality reduction and visualisation procedure for multi-omic data. Spectrum is provided as a R software package (https://cran.r-project.org/web/packages/Spectrum/index.html).

## Results

### Spectrum provides fast effective clustering of single and multi-omic data

First, we tested Spectrum’s ability to identify the ground truth K on individual simulated Gaussian datasets (Fig. S1). In each case, Spectrum correctly identified the optimal K. The method can also detect more complex non-Gaussian structures (Fig. S2). To demonstrate performance of Spectrum on real data from a single platform, we ran the algorithm on seven TCGA RNA-seq datasets^1–7^. We used log-rank tests to evaluate the significance of survival time differences between identified clusters. Comparison of Spectrum p values with those from CLEST, M3C, PINSplus, and SNF found that Spectrum performed substantially better overall in finding clusters significantly related to patient survival (Fig. 1a). For comparing different methods, we took both a rank and p value based approach to assess performance, individual p values and rankings for each method on each dataset are included in Supplementary Table 1. To give an initial indication of the relative computational resources required for a single platform analysis, algorithm runtime was investigated on a kidney cancer RNA-seq dataset^6^ with 240 samples and 5000 features. This analysis was performed on a single core of an Intel Core i7-6560U CPU @ 2.20GHz laptop computer with 16GB of DDR3 RAM. Spectrum was the fastest method (1.13 seconds), closely followed by SNF (2.67 seconds), PINSplus was still fast (8.53 seconds), M3C (123.91 seconds) and CLEST (283.34 seconds) were both considerably slower.

**Figure 1.**
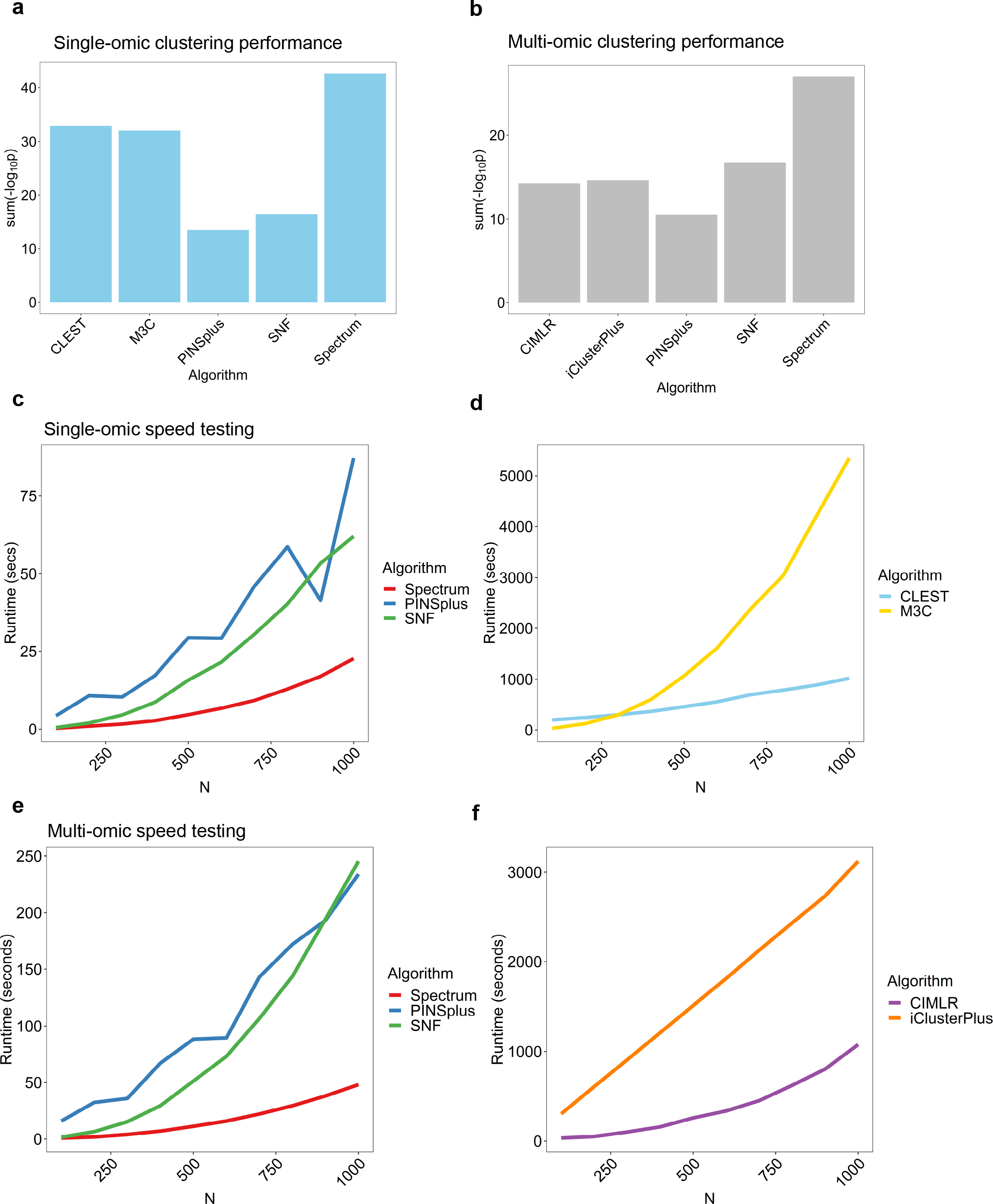
Spectrum provides fast and effective clustering for single and multi-omic TCGA data. The clustering performance score is the sum of the −log_10_(p values) from a Cox proportional hazards regression model using a log-rank test to assess significance across seven TCGA datasets (ranks are also included in Table I). (A) Spectrum relative performance just on RNA-seq datasets. (B) Spectrum relative performance on multi-omic datasets (mRNA, miRNA, protein). (C) Runtime analysis for faster single-omic clustering algorithms. (D) Runtime analysis for slower single-omic clustering algorithms. (E) Runtime analysis for faster multi-omic clustering algorithms. (F) Runtime analysis for slower multi-omic clustering algorithms.

**Table I.** Spectrum multi-omic clustering performance relative to other algorithms. P values are from a Cox proportional hazards regression model using a log-rank test to test the significance of the survival time differences between clusters. In brackets next to the p values are the ranks for each dataset. The first final row is the summed −log10(p values) for that column (higher is better), the second is the sum of the ranks (lower is better). PCPG stands for Pheochromocytoma and Paraganglioma. For all datasets, the three data types used were mRNA, miRNA, and protein.

| Dataset | N | Spectrum | PINSplus | iClusterPlus | SNF | CIMLR |
| --- | --- | --- | --- | --- | --- | --- |
| Bladder<sup>7</sup> | 338 | 0.0042 (3) | 0.31 (5) | 0.0022 (2) | 0.00022 (1) | 0.0047 (4) |
| Brain<sup>3</sup> | 425 | 3.76E-16 (1) | 0.0053 (4) | 1.72E-07 (3) | 4.17E-11 (2) | 0.013 (5) |
| Breast<sup>4</sup> | 634 | 1.47E-07 (1) | 2.85E-05 (4) | 1.78E-05 (3) | 0.94 (5) | 2.04E-07 (2) |
| Kidney<sup>6</sup> | 240 | 0.91 (5) | 0.038 (2) | 0.24 (4) | 0.045 (3) | 0.0026 (1) |
| PCPG<sup>5</sup> | 80 | 0.043 (1) | 0.18 (4) | 0.093 (3) | 0.09 (2) | 0.54 (5) |
| Skin<sup>2</sup> | 338 | 0.0014 (1) | 0.96 (5) | 0.4 (3) | 0.51 (4) | 0.0029 (2) |
| Thyroid<sup>1</sup> | 219 | 0.049 (1) | 0.09 (2) | 0.67 (5) | 0.18 (4) | 0.17 (3) |
| P value score |  | 30.21 | 10.56 | 16.40 | 17.49 | 17.06 |
| Rank score |  | 13 | 26 | 23 | 21 | 22 |

Next, we conducted a multi-omic data simulation that generated Gaussian clusters with added noise (Fig. S3a) to test the behaviour of Spectrum’s data integration system. Individual platform clustering using Spectrum did not detect the optimal K (Fig. S3b,c). However, using the TPG data integration and diffusion method, Spectrum identified the optimal K for the combined dataset (Fig. S3d). We proceeded to test Spectrum’s ability to detect clusters with significant differences in survival time on seven multi-omic TCGA datasets relative to other methods (Table 1). The analysis included mRNA, miRNA, and protein data. Similar to our observations on a single platform, Spectrum performed very well (Fig. 1b). Spectrum performed particularly well on the larger datasets with greater potential for clinical significance, namely, the breast (p = 1.47E-07) and brain cancer (p = 3.76E-16) datasets. Next, to gain an initial insight into relative multi-omic runtimes we tested the algorithms on the kidney TCGA dataset^6^. Spectrum performed the fastest (2.5 seconds), followed by SNF (4.06 seconds), PINSplus (27.22 seconds), CIMLR (59.56 seconds), and iClusterPlus (305.35 seconds). A more detailed analysis of runtime was performed for all single and multi-omic algorithms using simulated data. This worked by increasing the number of samples from 100 to 1000 in steps of 100, with each dataset containing 5000 features (Fig. 1c-f). These analyses demonstrated the greater speed of Spectrum relative to other methods. Spectrum’s strong performance in finding clinically related clusters comes with a bonus of faster runtimes.

We next demonstrated the advantage of Spectrum’s adaptive density-aware kernel by comparison with the classic Zelnik-Manor kernel^21^, a non-density-aware kernel that adapts to local data scale only. First, Spectrum using either of the two kernels was run on a non-Gaussian synthetic dataset consisting of two worm structures. The clustering demonstrated that the density-aware kernel improved the classification (Fig. S4a,b). Next differences on TCGA multi-omic data were examined. Analysis of the brain cancer multi-omic dataset^3^ found the density-aware kernel detected two additional clusters in comparison with the Zelnik-Manor kernel (Fig. 2a). Spectrum’s method for visualising multi-omic clusters was demonstrated. It runs Uniform Manifold Approximation and Projection (UMAP)^28^ on the integrated similarity matrix (Fig. 2a), this is a new method inspired by the recent approach applied in single-cell RNA-seq analysis by SIMLR^17^. Plots of the reduced data from UMAP demonstrated the density-aware kernel results in more compact clusters than the Zelnik-Manor kernel. This was expected, as the adaptive kernel strengthens local connections in the graph where the samples are in regions of high density.

**Figure 2.**
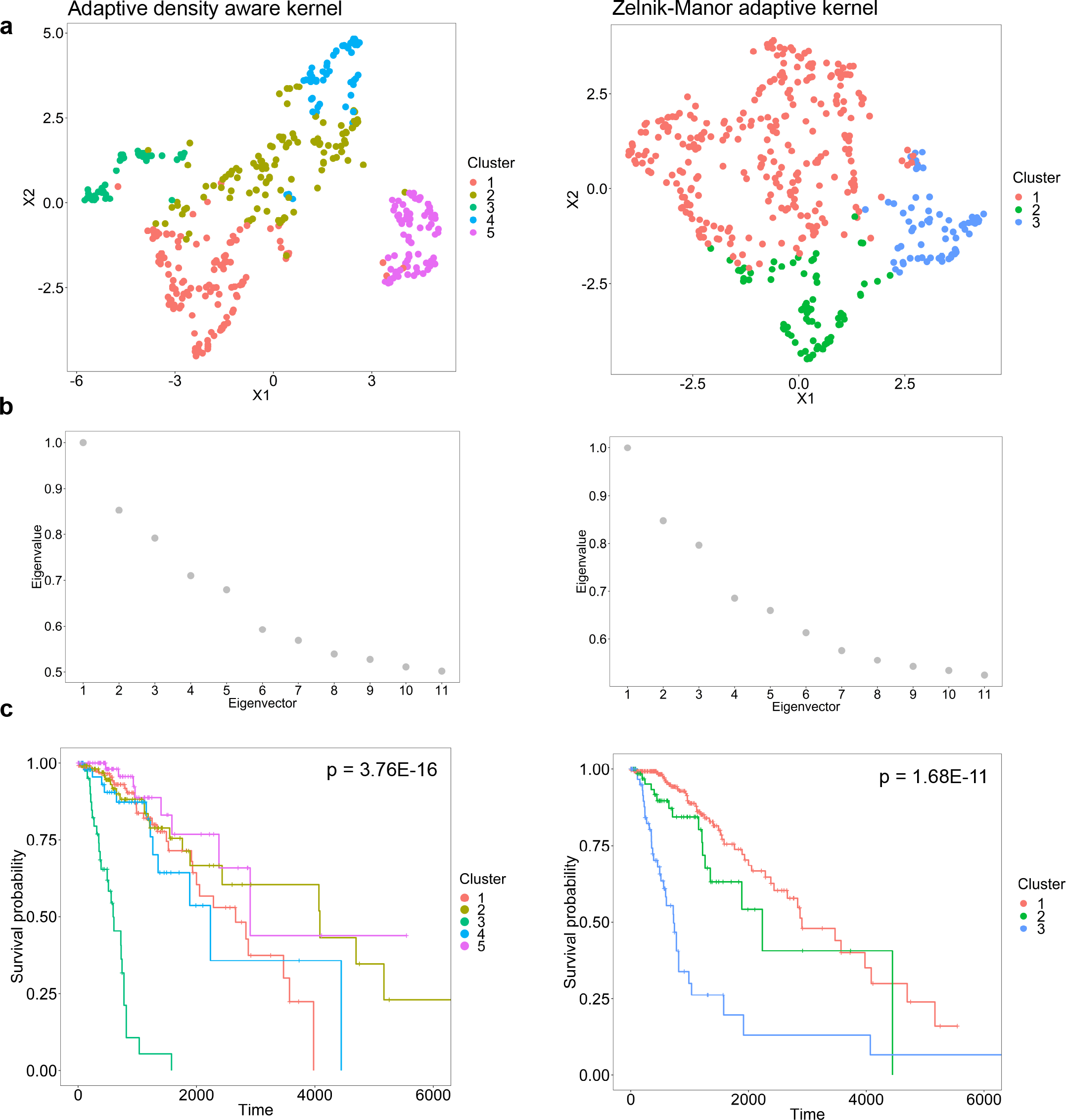
The adaptive density-aware kernel demonstrates an advantage in multi-omic analysis. On the right-hand side of the panel are the results for the Zelnik-Manor kernel^21^, while the density-aware kernel results are shown on the left-hand side. (A) Spectrum clustering assignments from the brain cancer dataset^3^, UMAP was run on the integrated similarity matrices for mRNA, miRNA, and protein data to generate the plots. (B) Eigenvalues for the eigenvectors of the graph Laplacians. The maximum eigengap was taken in each case as the optimal K. (C) Survival curves with p values from a Cox proportional hazards regression model using a log-rank test to assess significance between clusters.

The eigengap method Spectrum uses to decide on K by default was demonstrated (Fig. 2b). For the density-aware kernel, the greatest gap is between eigenvectors five and six, implying an optimal K of five, while for the Zelnik-Manor kernel, the greatest gap is found between eigenvectors three and four, implying an optimal K of three. The survival p values produced by the different methods were shown on survival curves (Fig. 2c). Spectrum obtained a greater level of significance using the density-aware kernel (p = 3.76E-16) than using the Zelnik-Manor kernel (p = 1.68E-11). We expanded this comparison to include all seven TCGA multi-omic datasets to find that the density-aware kernel has a noticeable advantage over the Zelnik-Manor non density-aware kernel (Table S2). These findings demonstrate the improvement gains by using a kernel that considers local density.

### Spectrum performs well at identifying cell types in single-cell RNA-seq data

We examined Spectrum’s performance on simulated datasets that resemble single-cell RNA-seq, as they were made to consist of many Gaussian blobs that can overlap (Fig. S5). Spectrum identified the correct K for both the K=10 simulated dataset and the K=20 dataset (Fig. S5a-d). We examined Spectrum’s performance relative to other methods on seven real single-cell RNA-seq datasets^24–30^ by comparing the assigned clusters with the provided cell type labels using Normalised Mutual Information (NMI) (Table 2). Spectrum achieved the highest summed NMI (NMI=5.89), this was closely followed by Seurat (NMI=5.74), MUDAN (NMI=5.71), SC3 (NMI=5.49), and SIMLR (NMI=5.01). Spectrum’s summed NMI was favourably weighted by its performance on the Pollen dataset^27^ (NMI = 0.95). Using a rank-based score to eliminate this advantage, Spectrum still came joint first with SC3 (Table 2). T-distributed stochastic neighbour embedding (t-SNE) plots showing the Spectrum clustering assignments were produced (Fig. 3a-f, Fig. S6a,b). Overall, Spectrum, Seurat, SC3, and MUDAN performed similarly in these comparisons, however, SIMLR did not perform as well (Table 2).

**Table II.** Spectrum single cell RNA-seq clustering performance relative to other algorithms. Values used for scoring each algorithm refer to Normalised Mutual Information (NMI) of given cell type labels versus those defined by the clustering algorithm. The bracketed values are the rank for that algorithm relative to the others for each dataset. The first final row corresponds to the summation of the columns NMI values, the second corresponds to the summation of the ranks.

| Dataset | Cells | Spectrum | Seurat | SIMLR | SC3 | MUDAN |
| --- | --- | --- | --- | --- | --- | --- |
| Camp<sup>25</sup> | 777 | 0.85 (3) | 0.87 (1) | 0.67 (4) | 0.51 (5) | 0.86 (2) |
| Li<sup>27</sup> | 561 | 0.88 (2) | 0.86 (3) | 0.5 (4) | 0.89 (1) | 0.86 (3) |
| Patel<sup>29</sup> | 430 | 0.76 (5) | 0.78 (4) | 0.79 (3) | 0.84 (1) | 0.81 (2) |
| Pollen<sup>30</sup> | 301 | 0.96 (1) | 0.86 (4) | 0.79 (5) | 0.89 (2) | 0.87 (3) |
| Darmanis<sup>26</sup> | 466 | 0.72 (3) | 0.78 (1) | 0.74 (2) | 0.74 (2) | 0.7 (4) |
| Baron<sup>24</sup> | 8569 | 0.84 (1) | 0.76 (3) | 0.79 (2) | 0.76 (3) | 0.79 (2) |
| Muraro<sup>28</sup> | 2126 | 0.88 (1) | 0.83 (3) | 0.73 (5) | 0.86 (2) | 0.82 (4) |
| NMI score |  | 5.89 | 5.74 | 5.01 | 5.49 | 5.71 |
| Rank NMI score |  | 16 | 19 | 25 | 16 | 20 |

**Figure 3.**
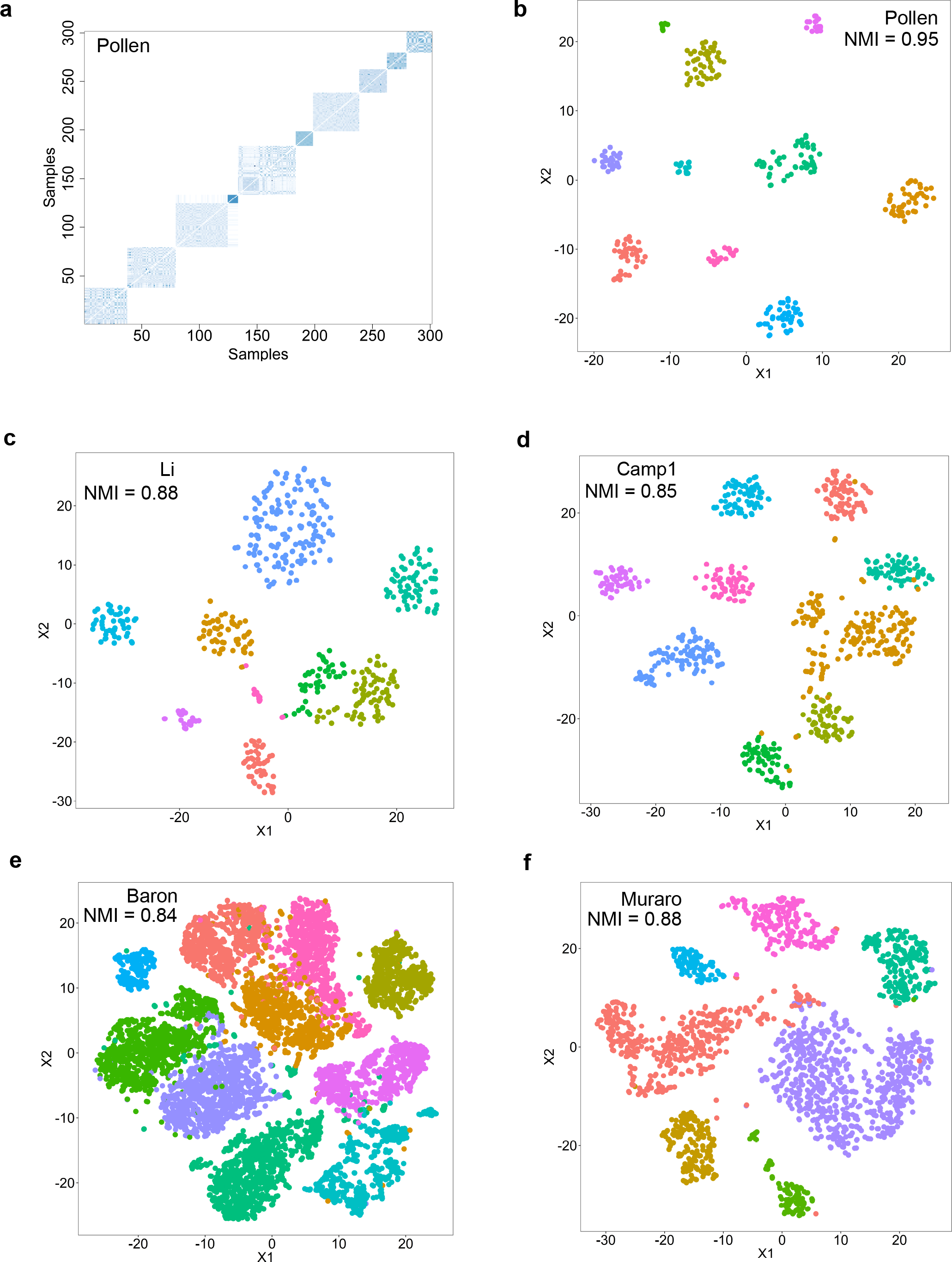
Spectrum performs well at identifying cell types in single cell RNA-seq data. For each dataset normalised mutual information (NMI) between the cell labels and the detected clusters is shown. (A) Heatmap of similarity matrix from Spectrum for the Pollen dataset^27^. (B) t-SNE of the Pollen data^27^ overlaid with clustering assignments. (C) Same as B, but from the Li data^25^. (D) Same as B, but from the Camp data^23^. (E) Same as B, but from the Baron data^24^. (F) Same as B, but from the Muraro data^28^.

In the described comparative analysis (Table 2), since the Baron^24^ and Muraro^28^ datasets had higher numbers of samples, to reduce runtime Spectrum was run using the FASP method (with 900 centroids). Even with the FASP data compression for these two datasets, Spectrum yielded the highest NMI relative to the other methods. Comparing Spectrum runtime on the Baron dataset (N=8569) yielded 1.95 hours without FASP versus 14.23 seconds with FASP, analyses were performed on a single core of an Intel Core i7-6560U CPU @ 2.20GHz laptop computer with 16GB of DDR3 RAM. On the Muraro dataset (N=2126), without FASP took 1.97 minutes and with took 11.83 seconds. Since the complexity of spectral clustering is cubic, *0*(*N*^3^) and the complexity of k means is linear, *0*(*KNT*), where *T* is the number of k means iterations, using k means as a precursor to compress the data (FASP) is computationally advantageous on larger datasets.

To gain an initial insight into relative runtimes of all methods (without using Spectrum’s FASP implementation) methods were run on the Camp dataset^23^ (777 samples). This analysis found MUDAN performed the fastest (0.23 seconds), followed by Seurat (2.45 seconds), Spectrum (12.64 seconds), SC3 (183.66 seconds), and SIMLR (264.31 seconds). A runtime analysis was performed for all algorithms on simulated datasets with 500 to 4000 samples in steps of 500 with 1000 features (Fig. S7). Spectrum was in the middle in terms of speed, usually performing faster than the SC3 algorithm. However, SC3 adjusted its own parameters to work faster at higher numbers of samples making it of comparable speed to Spectrum. Spectrum was slower than MUDAN and Seurat, but much faster than SIMLR. Overall, these data demonstrate Spectrum is well suited to clustering small to large single cell RNA-seq datasets, with FASP required for the later.

### A fast new heuristic for finding K when performing spectral clustering

Since the eigengap method does not automatically recognise both Gaussian and non-Gaussian structures (Fig. S1,2), we developed a complementary method which can. The method involves examining the multimodality of the eigenvectors of the data’s graph Laplacian, so we call it ‘the multimodality gap’. To demonstrate this method, five Gaussian blobs were generated (Fig. 4a) and the multimodality of the data’s graph’s eigenvectors were also displayed (Fig. 4b). The dip-test statistic^31^ (Z) which measures multimodality demonstrated a large gap between eigenvectors five and six. Therefore, using this method it was correctly concluded that K=5. Each individual eigenvector was plotted out (Fig. 4c) to visualise the changing distribution of the eigenvectors. As observed in the analysis of the set of Z values, there was a transition from a multimodal distribution at eigenvector five to a unimodal distribution at eigenvector six, confirming K=5. To further validate the method, several simulations were run and the method successfully clustered both complex non-Gaussian (Fig. S8a,b,c,d) and Gaussian clusters (Fig. S9a,b,c,d). However, since the simple method of looking for the greatest gap in the set of Z values can get stuck in local minima (Fig. S9d), the method was further enhanced by adding an algorithm to search for the last substantial gap.

**Figure 4.**
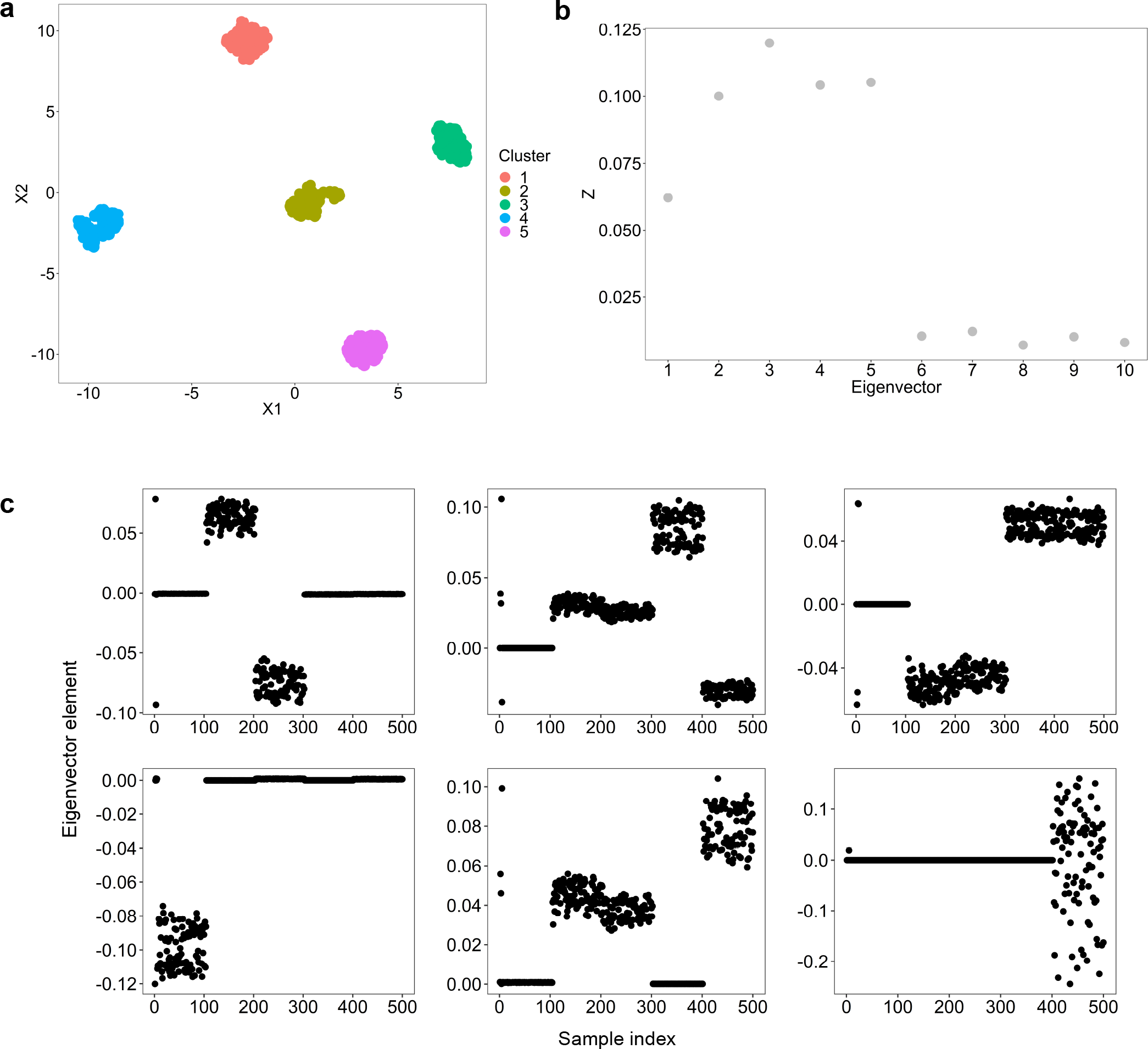
Demonstration of a new heuristic for finding K when spectral clustering. (A) PCA showing a synthetic dataset with five Gaussian clusters. (B) Using the multimodality drop method to find the ground truth K, the last substantial drop is in-between eigenvectors five and six, this refers to an optimal K of five (see methods). (C) Plots showing the individual eigenvectors of the graph Laplacian generated from the synthetic data. The distribution of eigenvector elements for the sixth eigenvector is far more unimodal than the fifth, demonstrating a solution of five is preferable.

We found the multimodality gap requires kernel tuning to perform well on certain datasets. This was evident in non-Gaussian data simulations, as with kernel tuning there is a perfect clustering result for the spirals test data (Fig. S10a), while without kernel tuning the method fails to cluster correctly (Fig. SlOb). Kernel tuning is performed by simply changing the *P* parameter of the self-tuning kernel and for each kernel finding the maximum multimodality gap between any pair of eigenvectors. The kernel that yields the greatest gap is the optimal kernel, where the most negative D value corresponds to that kernel with the maximum gap (Fig. S10c).

We examined the performance of the multimodality gap across the seven TCGA multi-omic datasets to demonstrate its applicability as a complementary method to the eigengap. This analysis found the multimodality gap can provide different p values compared with the eigengap (Table S3). Optimal methods will vary according to the data. For example, the multimodality gap (p = 0.0019) has a lower p value than the eigengap (p = 0.91) on the kidney cancer data^6^. Including a second method to automatically decide K gives the user power to find the best approach for their data. This method also presents a novel solution to an open problem in spectral clustering.

## Discussion

Spectrum provides fast, flexible spectral clustering for complex genome wide expression data. Previous tools have been developed for a single purpose, for example multi-omic clustering^12–14^, single-omic clustering^9,10^, or single cell RNA-seq clustering^15,17^. However, Spectrum has been demonstrated to work well on all these tasks and therefore is distinct to other methods. Our experiments demonstrated that Spectrum offers state-of-the-art performance in patient stratification, often identifying clusters with greater significant differences in survival time than other methods. Examining our density-aware kernel in comparison with the Zelnik-Manor kernel^21^ demonstrated Spectrum’s emphasis on strengthening local connections in the graph in regions of high density, partially accounts for its performance advantage. Spectrum is flexible and adapts to the data by using the k nearest neighbour distances instead of global parameters^18,22^ when performing kernel calculations.

Spectrum was the fastest method in single-platform and multi-omic TCGA analysis, this is partially due to code optimisation of the density-aware kernel calculation. Even though the kernel is relatively sophisticated, it runs quickly. However, single cell RNA-seq clustering tools are typically faster than single-omic clustering tools used on TCGA data. As we demonstrated, Spectrum performs well at single cell RNA-seq clustering, but has a drawback of increased runtime relative to the community detection methods Seurat and MUDAN. This is compensated for by implementation of a fast data compression method^23^ for massive datasets. Even without data compression, Spectrum performed similarly in speed to SC3 and much faster than SIMLR.

The new multimodality gap heuristic for finding K increases the flexibility of the method and increases the appeal of Spectrum to researchers in other disciplines. This is because it can recognise both complex shapes and Gaussian clusters. There are few good solutions to this problem, none of which are implemented in a publicly available R program. The Zelnik-Manor self-tuning algorithm^21^ involves a gradient descent technique that is complex to code and time consuming. In contrast, the multimodality gap is more straightforward, effective, and can be used to tune the kernel. Incorporating this method into Spectrum gives the user power to find the best approach for their data. Overall, Spectrum is well suited to clustering complex omic’ data and other data types. With its strong performance both in runtime and clustering results it should appeal to a broad range of data analysts.

## Methods

### Spectrum

Spectrum can be used to cluster continuous single or multi-omic data, the approach can also be used generally for single and multi-view clustering. Let us now describe the method. Let *E* denote an expression matrix *E* ∈ *R*^*N*×*M*^ where *N* is the number of samples and *M* is the number of features. If we are dealing with multiple datasets, then we have a set of *T* matrices *L* = {*E*_1_, *E*_2_, *E*_3_ …, *E*_*T*_}, where 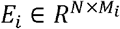 where *M*_*i*_ is the number of features per sample from the *i*^*th*^ data type. The first step of the algorithm is to calculate the similarity matrix or matrices using the adaptive density-aware kernel.

### Adaptive density-aware kernel

We first calculate the similarity matrix 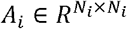 for the *i*^*th*^ expression matrix in *L*, where *N*_*i*_ equals the number of samples from the *i*^*th*^ data type. For the *i*^*th*^ data type, given a set of *N* points, 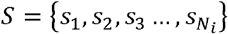, the adaptive density-aware kernel is:

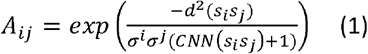

Where *d*(*s*_*i*_*s*_*j*_) denotes the Euclidean distance between points *s*_*i*_ and *s*_*j*_, *CNN*(*s*_*i*_*s*_*j*_) denotes the common nearest neighbours shared between the *i*^*th*^ and *j*^*th*^ samples in the *C*^*th*^ nearest neighbours neighbourhood of each sample. *σ*^*i*^ is the distance between the *i*^*th*^ sample and its *P*^*th*^ nearest neighbour. In the case of multiple datasets, this yields a new set of *T* similarity matrices, *Y* = {*A*_1_, *A*_2_, *A*_3_ …, *A*_*T*_}, while if we have just one platform, *T* = 1, we have a single similarity matrix *A*.

The adaptive density-aware kernel is inspired by the widely used self-tuning kernel by Zelnik-Manor^21^ and the more recent density-aware kernel by Zhang^22^. A drawback of the Zelnik-Manor kernel^21^, is that it does not adapt to the local density of the samples; however, it does adapt to scale using local *σ*^*i*^ and *σ*^*j*^ parameters. The problem with the Zhang kernel^22^ is that it falls back on the usage of a global *σ* parameter for the entire dataset, which is difficult and time consuming to tune. It also relies on a global radius parameter, *ε* that defines the density around the *i*^*th*^ and *j*^*th*^ samples when calculating the common nearest neighbours, *CNN*(*s*_*i*_*s*_*j*_). *ε*, like a global *σ*, is sensitive to the scale of the specific dataset. Therefore, our aim was to create a kernel that self-tunes to the scale and density of each new dataset by relying on the sample’s local statistics.

The adaptive density-aware kernel automatically tunes to the scale and density of each pair by examining the distance from each sample to their *P*^*th*^ nearest neighbour and their overlap within the *C*^*th*^ nearest neighbour neighbourhood of each sample. The *CNN*(*s*_*i*_*s*_*j*_) calculation tells us how many samples are shared between the *C*^*th*^ nearest neighbour neighbourhood of both samples. If these samples are close together in a region of high density, their connectivity will become stronger. Therefore, this kernel adds weight to locally dense connections. *C* and *P* are free parameters in this kernel. We set them to default values of 7 and 3, respectively. These parameters were used for all experiments in this manuscript, including those on simulated data, and perform well on a range of genome wide expression data.

### Combining multi-view data and tensor product graph diffusion

For combining the similarity matrices (graphs) in the set *Y*, Spectrum uses a recent technique from the machine learning literature^19^ that involves calculating a cross view tensor product graph (TPG) from each pair of individual graphs. The technique has so far not been applied in genome wide data analysis. Cross view TPGs capture higher order information of the data. The cross view TPGs are integrated using linear combinations to form a single graph. Graph diffusion is then performed to reveal the underlying data structure. Shu et al.,^19^ give a computationally efficient algorithm for this. The process is mathematically analogous to the TPG approach but can be calculated using a non TPG which makes the process much faster. Spectrum can also use a slight modification of this method for a single data type, more specifically:

1. Combine similarity matrices from the set *Y*. If we are dealing with a single similarity matrix, *T* = 1, then this step is skipped, but steps 2-5 are the same:

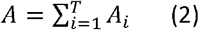
2. Sparsify *A* by keeping only the *Z*^*th*^ nearest neighbours of each sample *s*_*i*_ and setting the rest to 0. Let *R*_*i*_ be the set of *Z* nearest neighbour samples for *s*_*i*_. In this method we set *Z* = 10:

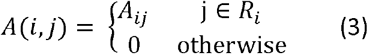
3. Row normalise *A*, so that each row sums to 1:

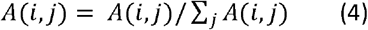
4. Perform graph diffusion iterations. Let *Q*^1^ = *A*, and *I* be the identity matrix for *A*. Then for the *t*^*th*^ iteration from 2… *iters*, where *iters* = 5:

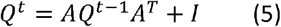
5. We then take the final similarity matrix as *A*\* = *Q*^*T*^. This ends the procedure. *A*\* can now be used as the input for spectral clustering.

The parameters *Z* = 10 and *iters* = 5 are set in alignment with previous work^19^.

### Spectral clustering of similarity matrix

Starting with *A*\*, Spectrum uses the Ng spectral clustering method^18^, but with the eigengap heuristic to estimate the number of clusters and Gaussian Mixture Modelling (GMM) to cluster the final eigenvector matrix. More specifically:

1. Using *D*, the diagonal matrix whose (*i, i*) element is the sum of *A*\*’s *i*^*th*^ row, construct the normalised graph Laplacian *L*:

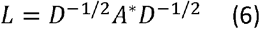
2. Perform the eigendecom position of *L* and thus extract its eigenvectors *x*_1_, *x*_2_, … *x*_*N*_ and eigenvalues λ_1_, λ_2_ …λ_*N*+1_.
3. Evaluate the eigengap for eigenvalues, starting with the second eigenvalue, *n* = 2, and choose the optimal k, denoted by *k*\*, for which the eigengap is maximised:

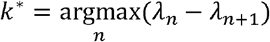
4. Get the *x*_1_, *x*_2_,… *x*_*k*\*_, *k*\* largest eigenvectors of *L*, then form the matrix, *X* = [*x*_1_, *x*_2_,… *x*_*k*\*_] ∈ ℝ^*N*×*k*\*^ by stacking the eigenvectors in columns.
5. Form the matrix *Y* from *X* by renormalizing each of *X*’s rows to have unit length:

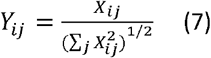
6. Now each row of *Y* is treated as a sample, *s*_*i*_ then all samples are clustered into *k*\* clusters using GMM. Spectrum uses the implementation of GMM from the ClusterR CRAN package.

### A heuristic for finding K that analyses eigenvector distributions

How to select the optimal K is an open question in the field of spectral clustering. A natural way to solve this problem when spectral clustering is analysis of the eigendecomposition of the graph Laplacian. The classical eigengap method is effective for Gaussian clusters, however, its rule must be modified to detect non-Gaussian structures thus limiting its applicability (Figure S2). We found a new heuristic for finding K that could be used for Gaussian or non-Gaussian structures and as a complementary method to analyse genome wide expression datasets. The method examines the multimodality of the eigenvectors of the graph Laplacian and looks for a point beyond which there is no more substantial decrease in multimodality.

Intuitively, the degree of multimodality defines how informative a given eigenvector is, and when we pass the optimal K moving along the sorted eigenvectors, *V* = {*x*_1_, *x*_2_, … *x*_*N*_}, we expect a large drop in useful information. Multimodality is quantified using the well-known dip test statistic^31^. The method works better if the nearest neighbour parameter of the kernel is tuned from *P* = 1,…,10. This is done by selecting the kernel that gives the maximum multimodality gap. Analysing the multimodality drop was inspired by work by Xiang et al.,^20^ where the authors select out the most informative eigenvectors using an Expectation Maximisation technique, then use GMM and the BIC to choose K. However, in our experiments, we found this method to be unreliable. We now detail an alternative procedure for finding K and tuning the kernel based on the analysis of drops in eigenvector multimodality.

Let the set of dip test statistics be *Z* = {*z*_1_, *z*_2_, *z*_3_ …, *z*_*N*_}, calculated from the eigenvectors, *V* = {*x*_1_, *x*_2_, … *x*_*N*_}. Note that larger values of *z*_*i*_ correspond to greater eigenvector multimodality. To calculate the multimodality difference between consecutive values, we use, *d*_*i*_ = *z*_*i*_ − *Z*_*i*–1_. Since we require two values to get *d*_*i*_, the calculation must begin at *i* = 2, which corresponds to the first pair of eigenvectors. Let the set of *d*_*i*_ values calculated from *Z* be, *D* = {*d*_1_, *d*_2_, *d*_3_ …,*d*_*maxK*–1_}, the steps for this are as follows:

1. Find the optimal kernel, *A*\*. Each kernel is calculated using equation 1 and the nearest neighbour parameter *P* is tuned via a search from *P* = 1,…, 10. To do this, calculate the *P*^*th*^ graph Laplacian (equation 6) from the *P*^*th*^ kernel. Obtain the eigenvectors of these, *V*_*P*_. Calculate *Z*_*P*_ from these eigenvectors, then *D*_*P*_. Get min (*D*_*P*_) for each *P*, yielding a set, *D*_*min*_ = {*d*_1_, *d*_2_, … *d*_10_}. *A*\* is the kernel that corresponds to *min*{*D*_*min*_}.
2. Get the KNN graph of *A*\*, row normalise, then perform diffusion iterations (equations 3–5).
3. Calculate the graph Laplacian, *L* (equation 6).
4. Perform eigendecomposition of *L* yielding the eigenvectors, *V* = {*x*_1_, *x*_2_,… *x*_*N*_}.
5. Calculate the dip test statistics *Z* for the eigenvectors in *V*, then calculate the differences of these, *D*.
6. Pass *D* into the algorithm described below for finding the last substantial drop in multimodality. Let *k*\* be the optimal K found by this method.
7. Continue with equations 4–6 corresponding to the Ng spectral clustering method, with GMM to cluster the final eigenvector matrix, with K set to *k*\*.

One could select K using the maximum multimodality eigengap. However, we found that this simpler method is susceptible to getting stuck in local minima (Fig. S6). This naturally led to making an algorithm to find the last substantial drop. For finding *k*\* from set *Z*, we now describe this straightforward algorithm that reads along the elements of *Z*, to find a point where there is no more substantial decrease in multimodality.

### Finding last substantial multimodality gap

This algorithm will search *D*, storing in memory the biggest difference in multimodality. Let that be *d*_*min*_. A more negative *d*_*i*_ corresponds a bigger drop in the elements of *Z*. The method then examines if there are any more points ahead of this drop (up until < *c*_*max*_ points) that are 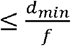, where *f* is the minimum magnitude that the drop must be to replace *d*_*min*_. If a new *d*_*min*_ is found, this new difference is stored in memory. This process continues until no more substantial drops are found with the threshold *c*_*max*_ to stop the search. More specifically:

1. Skip the first element of *D*, *d*_1_, as this corresponds to the drop from the 1^st^ to 2^nd^ eigenvector which is non-informative. Then, store in memory *d*_2_ as the greatest drop by default. Call this *d*_*min*_. Initialise a counter to *c* = 1, for keeping count of how many indices ahead we are of the stored *d*_*min*_.
2. Iterate from *d*_3_ … *d*_*maxK*–1_ and with each iteration check if 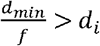. If so let *d*_*i*_ be the new *d*_*min*_, otherwise continue. If *c* > *c*_*max*_, break the loop and accept the current stored *d*_*min*_ as the solution.
3. The optimal number of classes is *k*\* = *i* − 1, where *i* is the index of *D* corresponding to the *d*_*i*_ taken as *d*_*min*_ in step 2.

The parameters used in this study for the multimodality drop procedure were *c*_*max*_ = 7 and *f* = 2, values we empirically selected based on our experience.

## Generating simulated datasets for analysis

Gaussian cluster simulations were all performed using the CRAN clusterlab package^9^, following the standard operating procedure. In the case of non-Gaussian structures, found throughout the Supplementary Figures, either the CRAN mlbench or clusterSim packages were used to simulate the data, using the default settings.

## Downloading and processing of real data for analysis

### TCGA datasets

The seven multi-omic TCGA datasets^1–7^ were downloaded from the Broad Institute (http://gdac.broadinstitute.org/). Pre-normalised data was used for each platform (mRNA, miRNA, and protein) and for every dataset each one was filtered in the same manner, using the coefficient of variation (CV) to select the top 50% most variable features. The data was then log _2_ transformed to reduce the influence of extreme values. Code for data pre-processing is found in the following GitHub repository (https://github.com/crj32/spectrum_manuscript). The processed multi-omic data is in Supplementary File 1. RNA-seq datasets were taken from the same studies as the multi-omic data and filtering of features was done in the same manner. However, more patients were included in the RNA-seq analyses because we did not have to unify the patient IDs between platforms. The RNA-seq data is included in Supplementary File 2. Code for performing log-rank tests in also in the github as well as commands for running methods.

### Single-cell RNA-seq datasets

The seven single-cell RNA-seq datasets^24–30^ were obtained from the Hemberg lab website (https://hemberg-lab.github.io/scRNA.sea.datasets/). For each dataset we used log_2_ normalised counts and selected the top 100 most variable genes for analysis. Code for data pre-processing is found in the manuscript’s GitHub repository (https://github.com/crj32/spectrum_manuscript). Additionally, we include the data ready for analysis in Supplementary File 3. Code for calculating NMI is in the github as well as commands for running methods.

## Author contributions

C.R.J conceived and designed the approach. C.R.J and D.W wrote the manuscript. C.R.J and D.W wrote the code. C.R.J performed data analyses. All authors reviewed and edited the manuscript. M.B, C.P, and M.L supervised the project.

## Additional information

The authors declare that they have no competing interests.

## Supplemental figure legends

**Figure S1. Spectrum performs well at identifying synthetic Gaussian clusters using eigenvalues.** In each case a Principal Component Analysis (PCA) of the synthetic data is shown alongside the eigenvalues of the eigenvectors of the data’s graph Laplacian. (A) Results from synthetic data analysis with K=2. (B) Analysis for K=3. (C) Analysis for K=4. (D) Analysis for K=5. In all cases the maximum eigengap method identified the correct number of clusters.

**Figure S2. Spectrum performs well at identifying non-Gaussian clusters using eigenvalues with a modified decision rule.** In each case a PCA of the synthetic data is shown alongside the eigenvalues of the eigenvectors of the data’s graph Laplacian. (A) Results from synthetic data analysis for K=2. (B) Analysis for K=3. In both cases the first non-zero eigengap method identified the correct number of clusters.

**Figure S3. Spectrum performs well at reducing noise when clustering data generated in a multi-omic data simulation.** (A) Three Gaussian clusters are simulated, each sample belongs to the same cluster in each simulation, but a degree of variability has been added. (B) Individual platform clustering could not detect the optimal K in every case due to noise. (C) Cluster assignments shown in PCAs using individual platforms with no data integration. (D) Spectrum data integration method results in finding K=3 on the combined data.

**Figure S4. The locally adaptive density-aware kernel outperforms the Zelnik Manor kernel on a non-Gaussian dataset.** (A) Spectrum results with the adaptive density-aware kernel. (B) Spectrum results with the Zelnik Manor non density-aware kernel.

**Figure S5. Spectrum performs well on simulated Gaussian data resembling single-cell RNA-seq.** (A) PCA of the K=10 synthetic dataset. (B) Eigenvalues of the eigenvectors of the data’s graph Laplacian, a maximum drop is located between 10 and 11 correctly designating this as the optimal K decision by the algorithm. (C) PCA of the K=20 synthetic dataset. (D) Eigengap method confirms K=20.

**Figure S6. Spectrum results from clustering the Patel single cell RNA-seq dataset.** (A) Results from t-SNE analysis of the Patel data^29^ overlaid with Spectrum clustering assignments. Normalised mutual information (NMI) between the cell labels and the detected clusters is shown. (B) Same as A, but for the Darmanis dataset^26^.

**Figure S7. Runtime analysis for single-cell RNA-seq clustering methods.** These analyses were run on a single core of an Intel Core i7-6560U CPU @ 2.20GHz laptop computer with 16GB of DDR3 RAM. Simulated datasets had 1000 features and varying number of samples (N). Spectrum was run without data compression (FASP).

**Figure S8. Demonstration of the multimodality gap method for recognising non-Gaussian structures.** Z refers to the dip test statistic *z*_*i*_ for the ith eigenvector, where a larger value indicates stronger multimodality. (A) Results using multimodality gap method on a two concentric circle dataset. The optimal K was identified because the gap between two and three is the last substantial one (see methods). (B) Results from analysing a three concentric circle dataset, again the correct K was identified. (C) Results from analysing a two half moon dataset, the correct K was detected. (D) Results from analysing a four cluster smiley face, the correct K was also found.

**Figure S9. Demonstration of the multimodality gap method for recognising Gaussian clusters.** Z refers to the dip test statistic *z*_*i*_ for the ith eigenvector, where a larger value indicates stronger multimodality. (A) Results from using the multimodality gap method on a K=2 Gaussian synthetic dataset. The optimal K of two is identified by the method because the eigenvalue gap between eigenvectors two and three is the last substantial one. (B) Results from using the multimodality gap method on a K=3 Gaussian synthetic dataset, the correct K is identified. (C) Results from using the multimodality gap method on a K=4 Gaussian synthetic dataset, the correct K is identified. (D) Results from using the multimodality gap on a K=5 Gaussian synthetic dataset, if we took the maximum multimodality gap the method fails, therefore we improved the method to find ‘the last substantial gap’. This is found by a greedy algorithm described in the methods.

**Figure S10. Kernel tuning can improve the multimodality gap method.** Z refers to *z*_*i*_ the dip test statistic for the ith eigenvector, where a larger value indicates stronger multimodality. (A) Results from the multimodality Spectral clustering method on spirals with kernel tuning of the *K* parameter that defines the number of nearest neighbours to use when calculating the local sigma (see methods). The correct K is found, as the last substantial gap is in-between eigenvectors two and three. (B) Same dataset, but the kernel tuning was not performed, and the correct K was not found. (C) Tuning results from changing the *K* parameter (NN). D refers to *D*_*min*_ the biggest difference between any consecutive elements of z in the set of dip statistics *Z* for a given value of *K*. A lower D indicates a bigger difference found for a given *K* and kernel, therefore indicating a preferred kernel.

