## Supplementary information for "Spectrum: Fast density-aware spectral clustering for single and multi-omic data"

### Supplementary figure 1

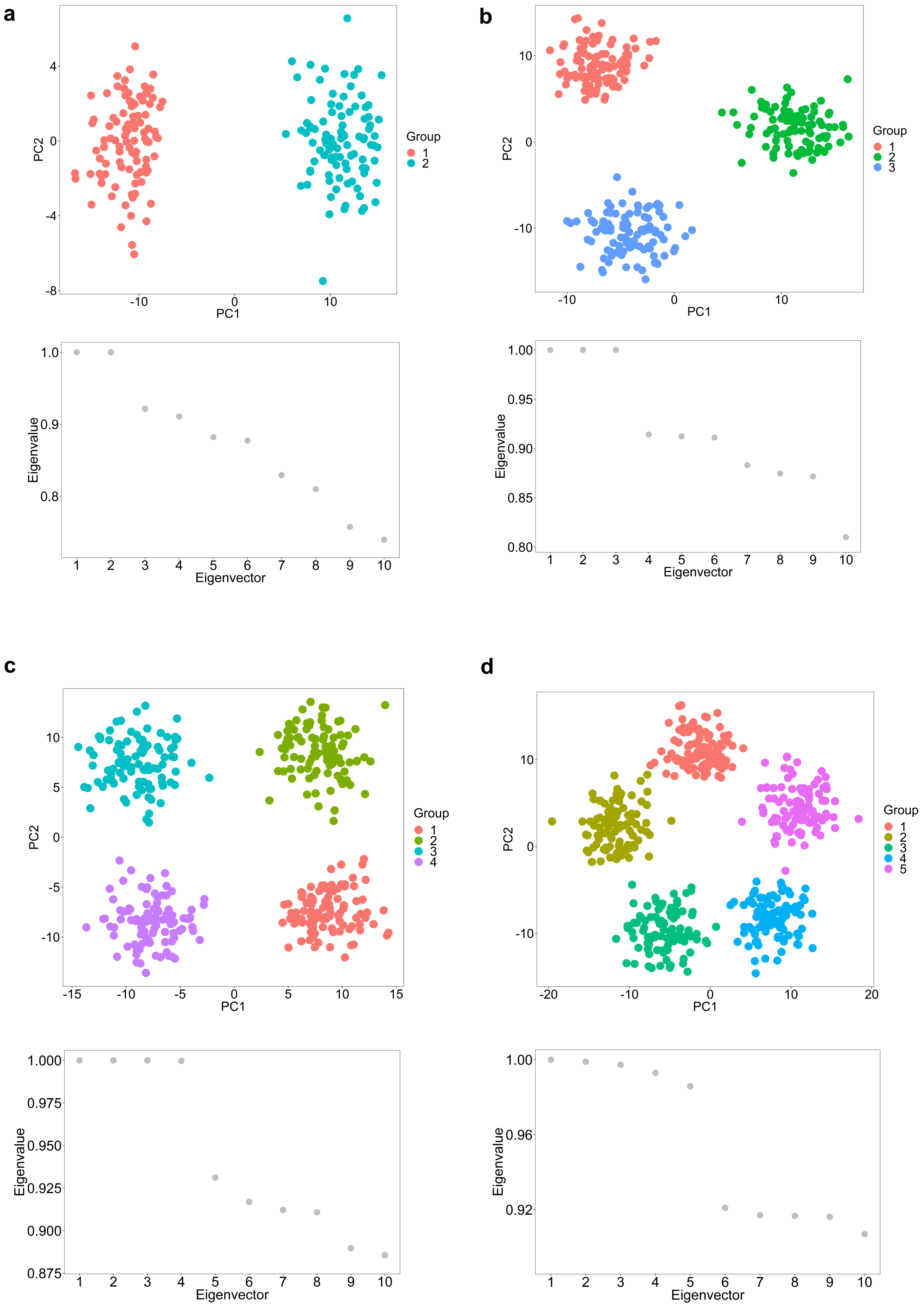

### Supplementary figure 2

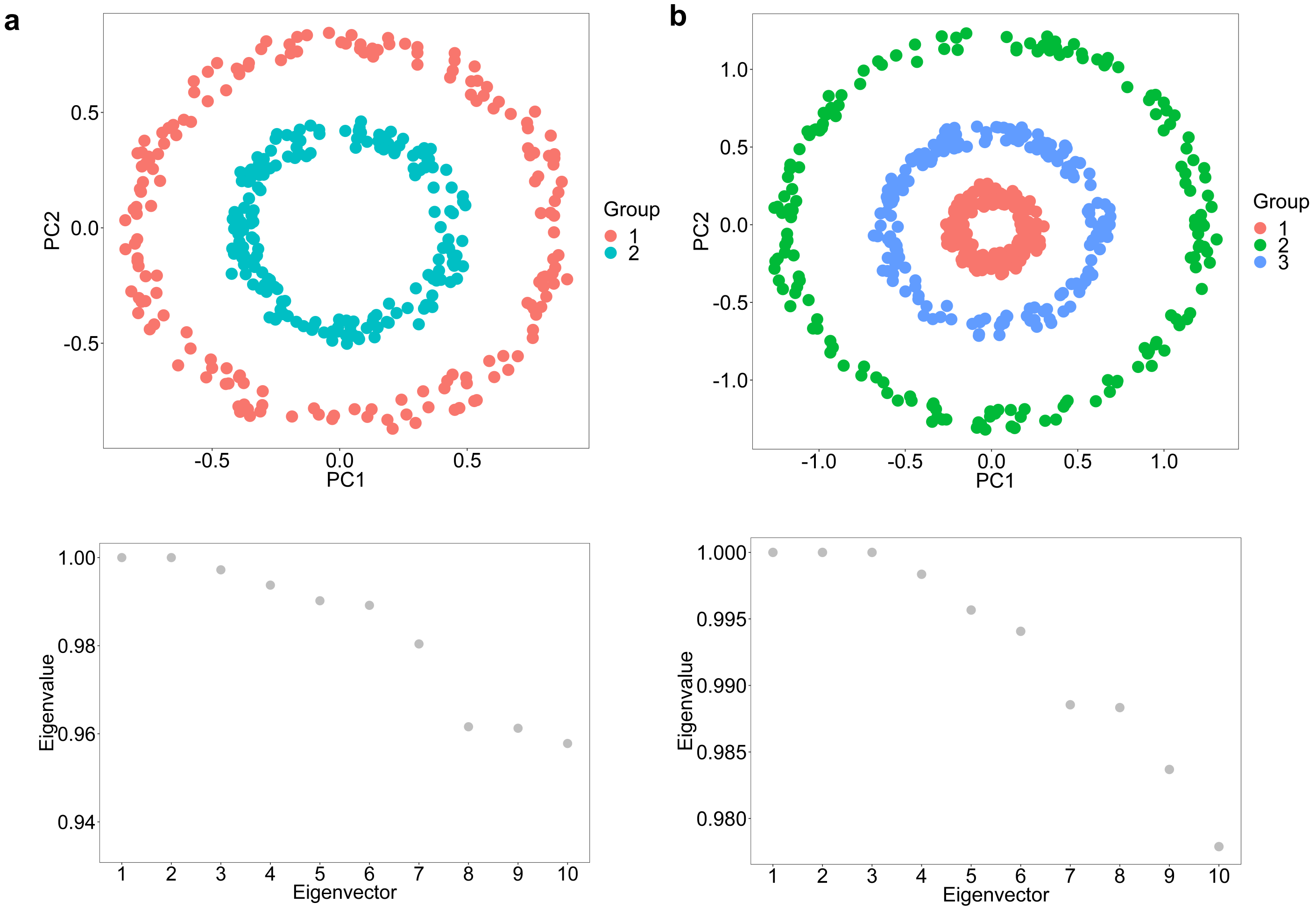

#### Supplementary figure 3

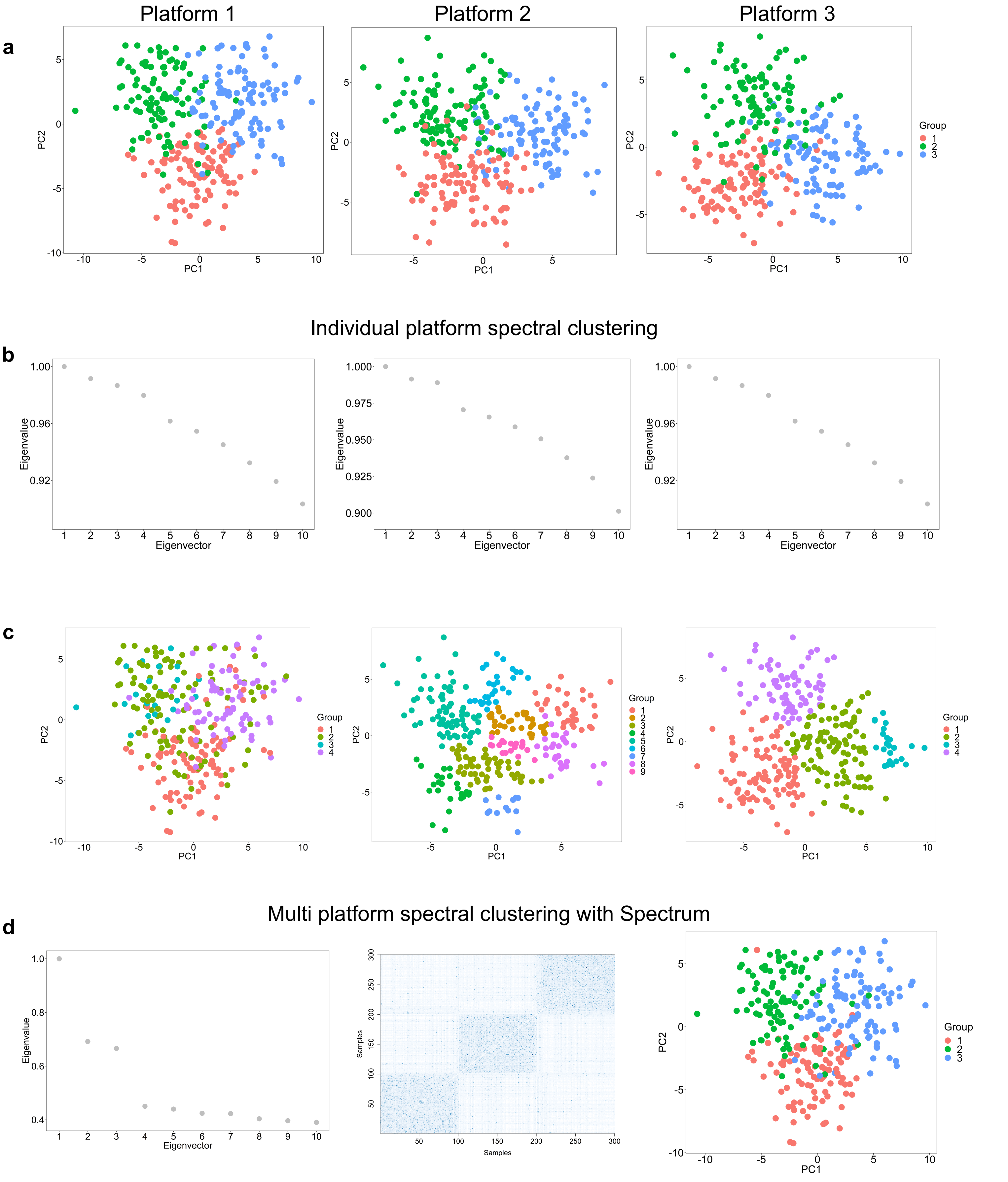

**Supplementary figure 4**

**a**

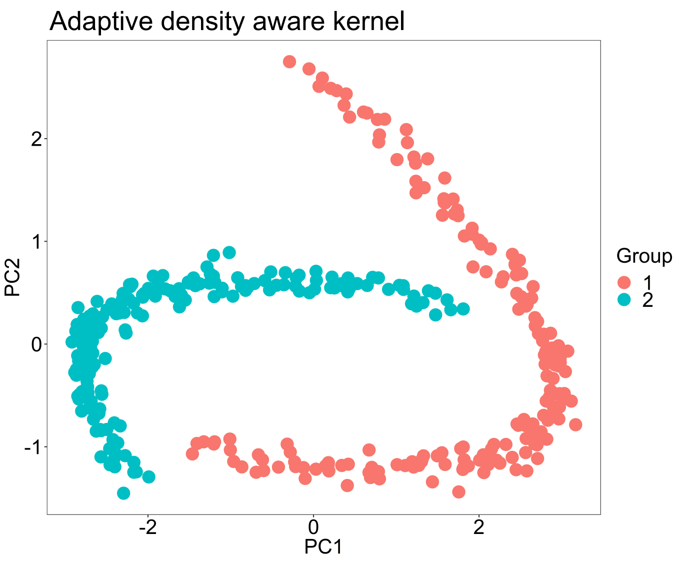


**b**

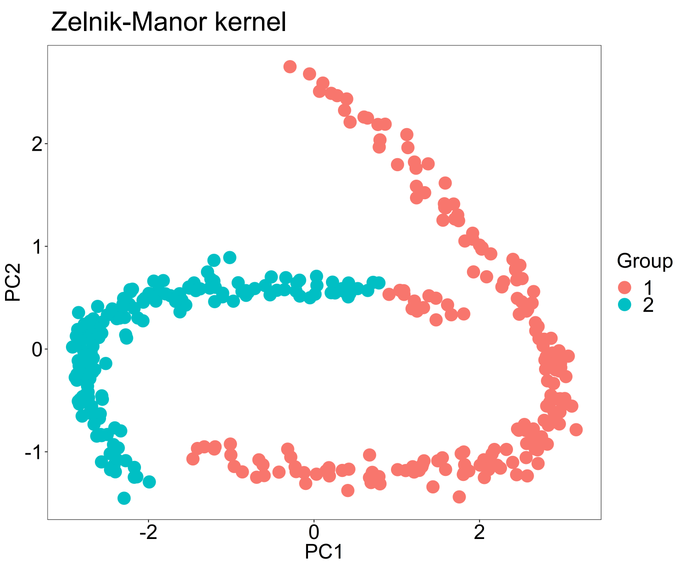

### Supplementary figure 5

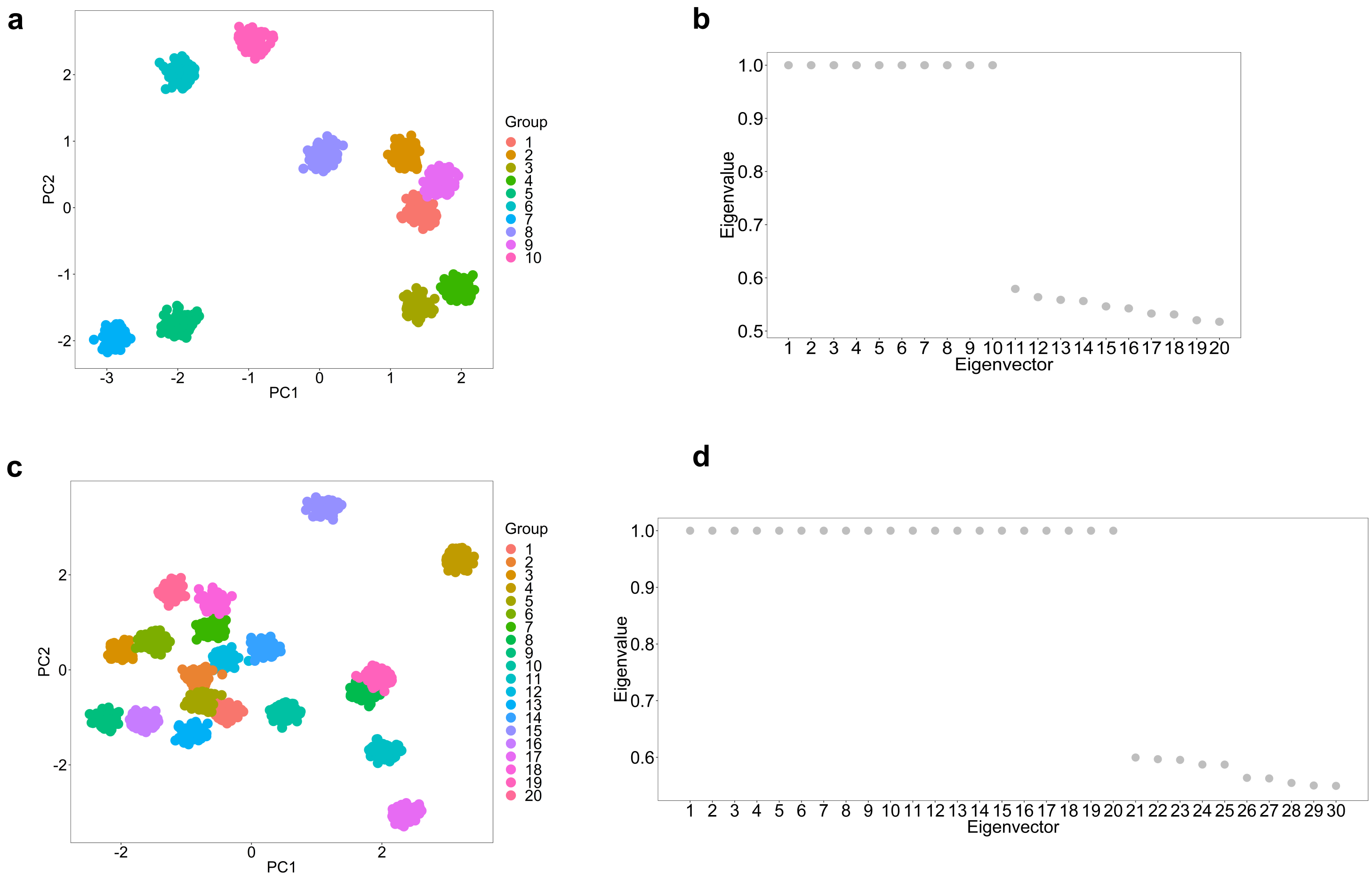

**Supplementary figure 6**

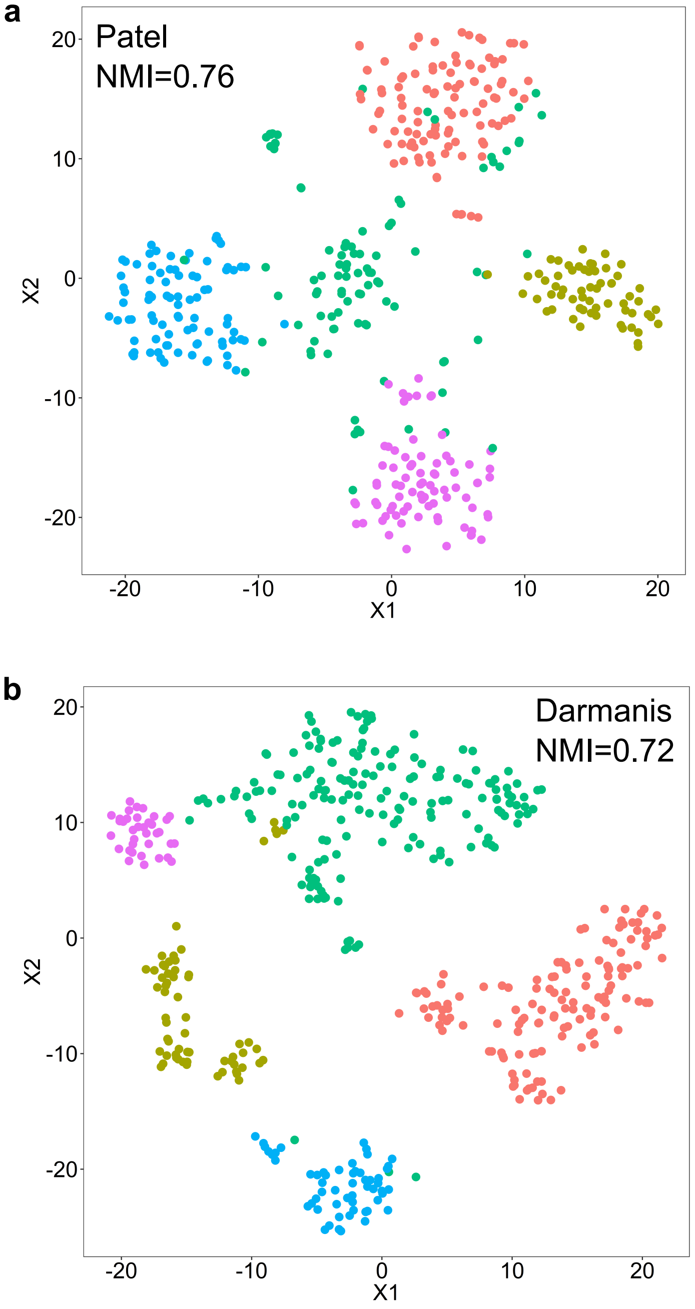

#### Supplementary figure 7

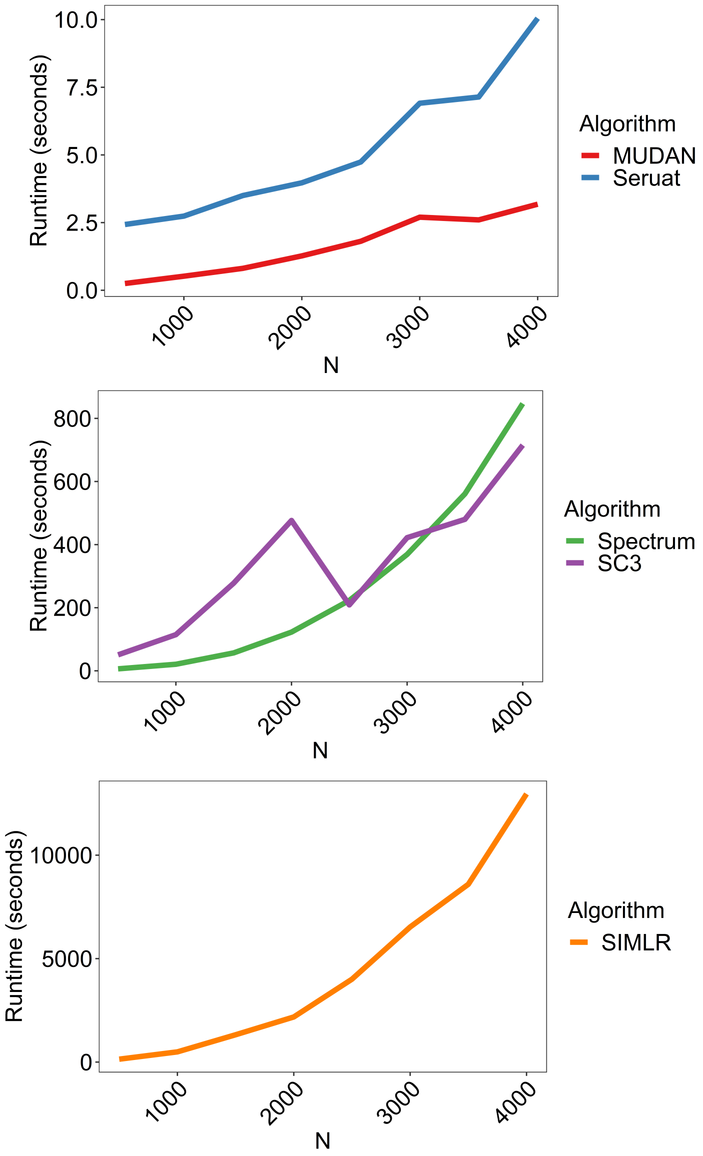

### Supplementary figure 8

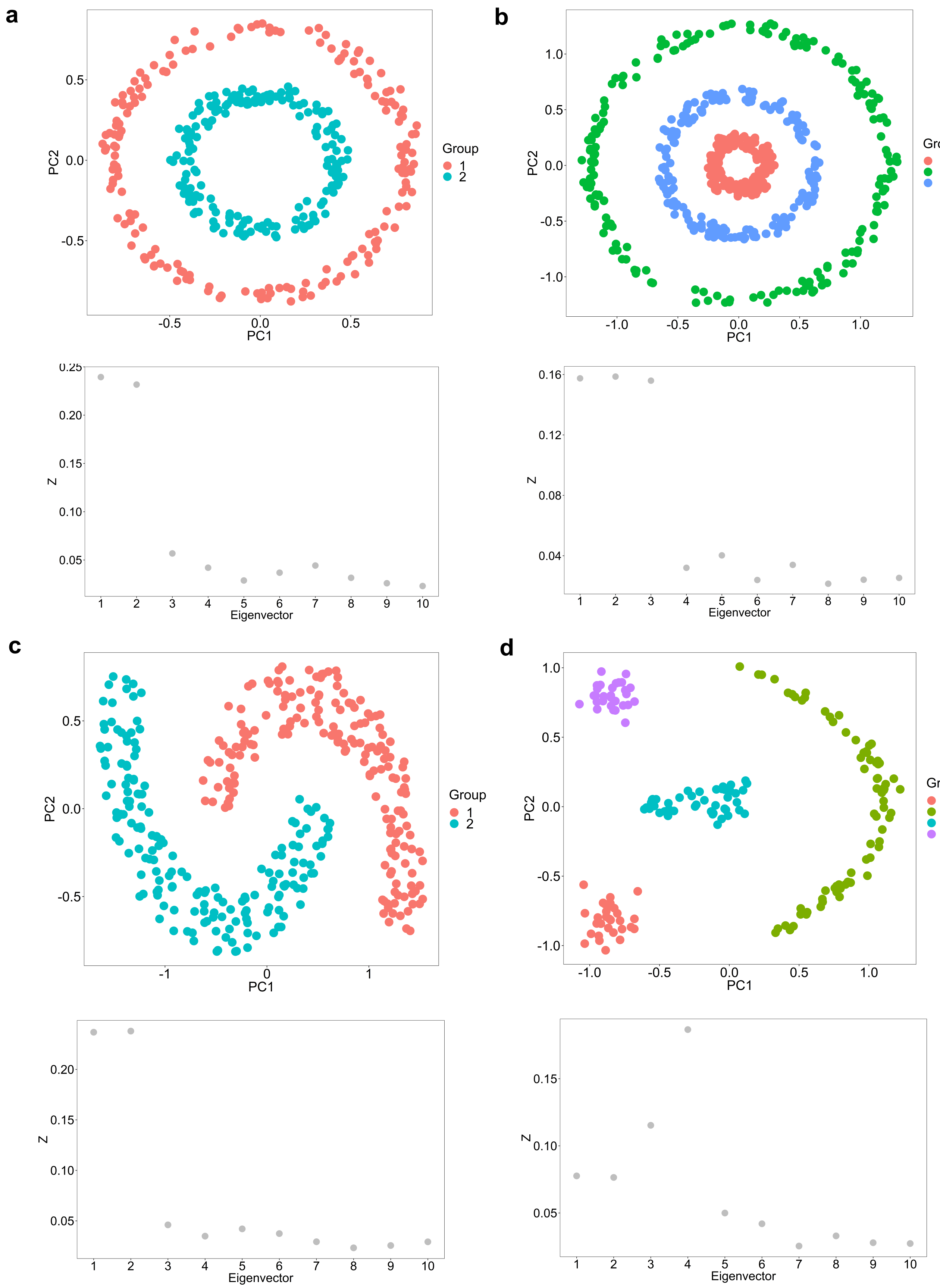

### Supplementary figure 9

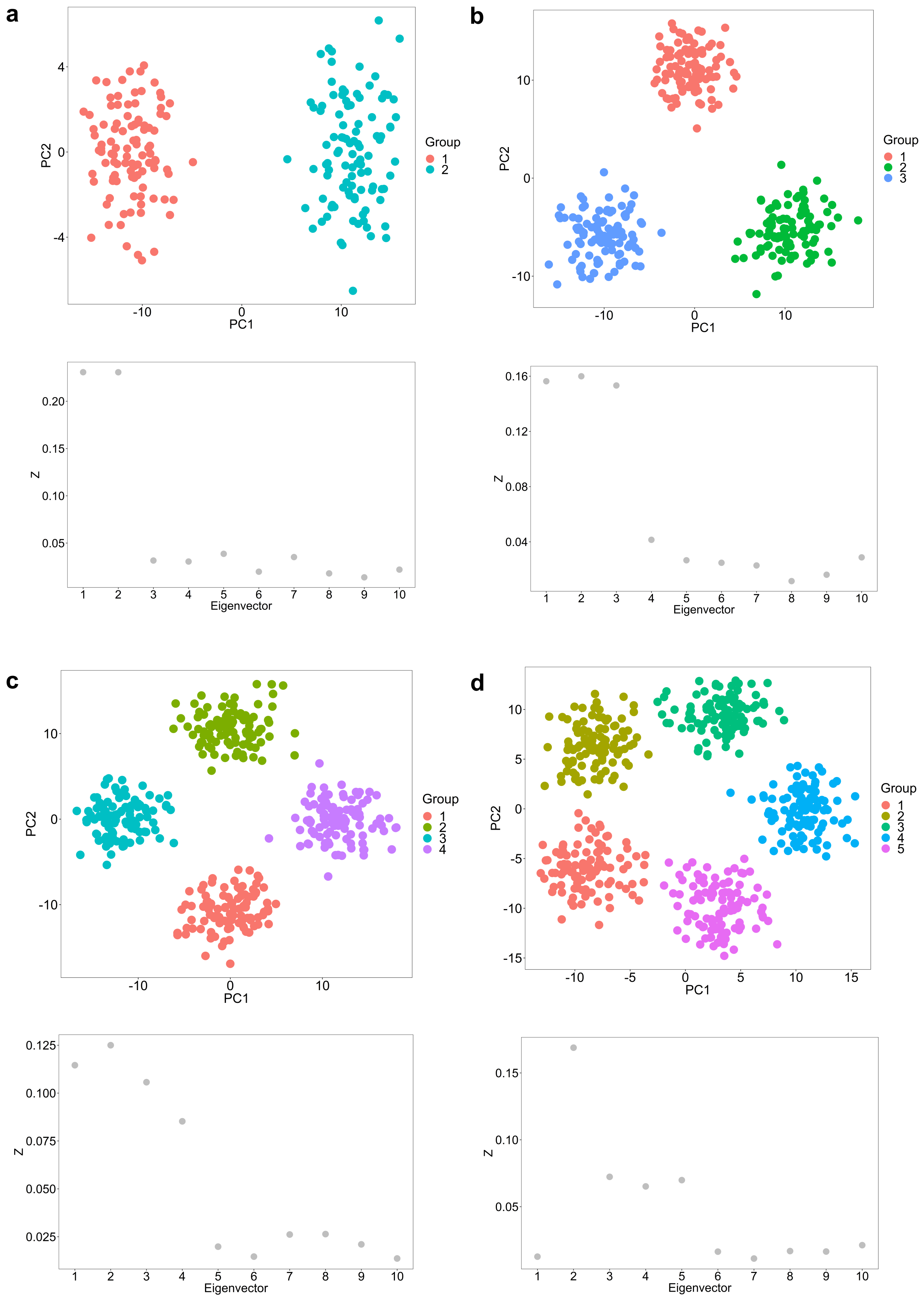

**Supplementary figure 10**

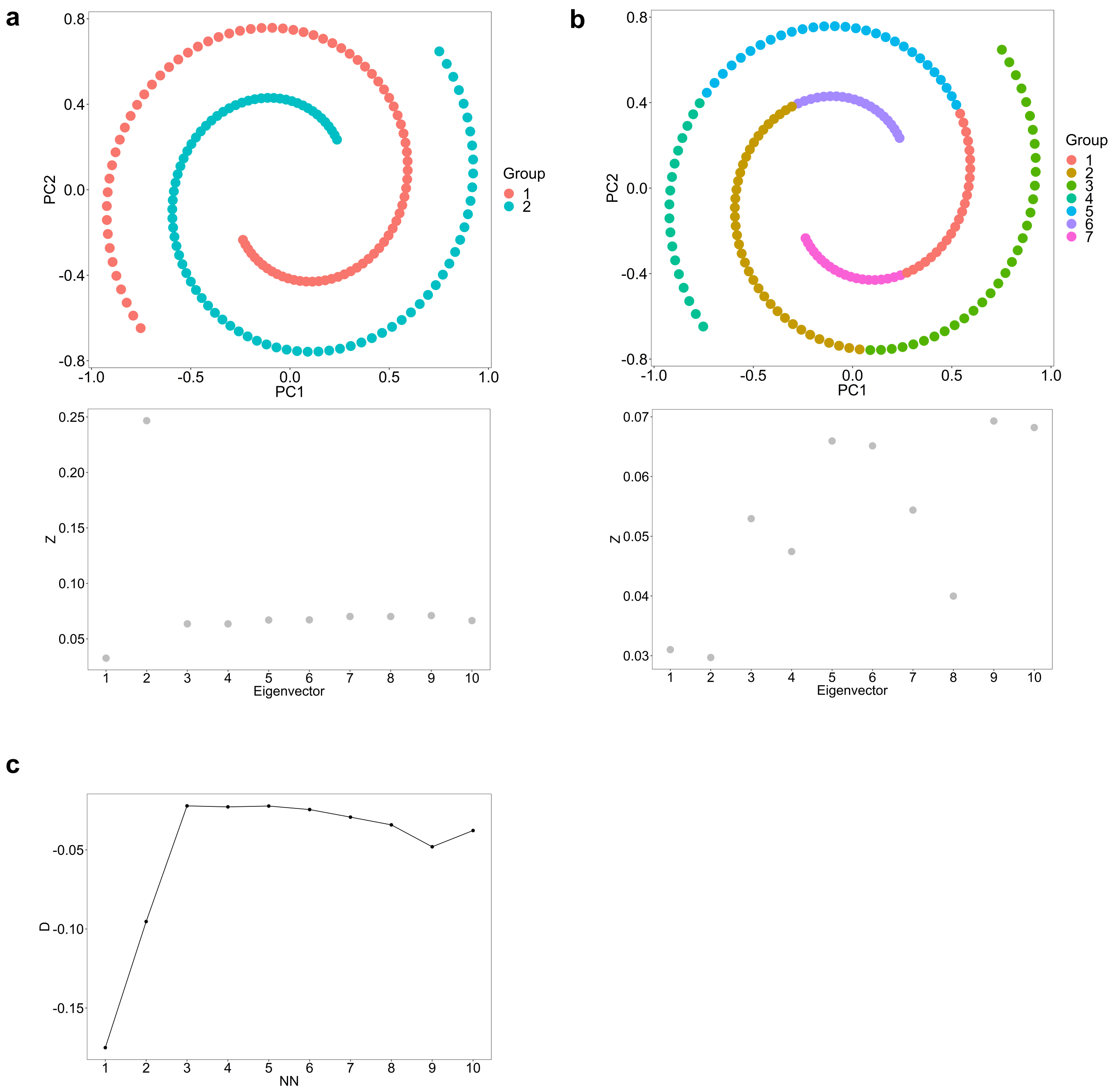

**Supplementary table I. Spectrum TCGA RNA-seq clustering performance compared with other algorithms.** P values are from a Cox proportional hazards regression model using a log-rank test to test the significance of the survival time differences between clusters. In brackets next to the p values are the ranks for each dataset (lowest p value corresponds to rank 1). The first final row is the summed -log10(p values) for that column (higher is better), the second in the sum of the ranks (lower is better). PCPG stands for Pheochromocytoma and Paraganglioma.

| Dataset | Datatype | N | Spectrum | PINSplus | M3C | SNF | CLEST |
|---|---|---|---|---|---|---|---|
| Bladder[7] | mRNA | 408 | 1.29E-05 (1) | 0.0013 (3) | 0.018 (4) | 0.00018 (2) | 0.0013 (3) |
| Brain[3] | mRNA | 515 | 4.03E-22 (1) | 0.46 (5) | 4.48E-16 (2) | 0.038 (4) | 4.71E-14 (3) |
| Breast[4] | mRNA | 1093 | 1.77E-05 (3) | 2.60E-06 (2) | 9.98E-07 (1) | 0.00033 (4) | 0.00035 (5) |
| Kidney[6] | mRNA | 533 | 1.98E-07 (1) | 0.4 (5) | 0.33 (4) | 0.3 (3) | 1.34E-06 (2) |
| PCPG[5] | mRNA | 179 | 0.4 | 0.56 | 0.36 | 0.46 | 0.43 |
| Skin[2] | mRNA | 469 | 0.00014 (4) | 0.00076 (5) | 1.14E-07 (1) | 8.70E-07 (2) | 9.31E-07 (3) |
| Thyroid[1] | mRNA | 501 | 0.21 | 0.11 | 0.084 | 0.14 | 0.11 |
| P value score | | | 42.67 | 13.54 | 32.04 | 16.42 | 32.90 |
| Rank score | | | 10 | 20 | 12 | 15 | 16 |

**Supplementary table II. Comparison of Spectrum density aware kernel versus the Zelnik-Manor et al. (2005) kernel in a multi-omic cluster analysis.** The final row corresponds to the sum of the $-\log_{10}$(p values) of that column.

| Dataset | Data types | Samples | Spectrum - density aware | Spectrum - Zelnik-Manor |
|---|---|---|---|---|
| Bladder[7] | mRNA, miRNA, protein | 338 | 0.0042 | 0.0033 |
| Brain[3] | mRNA, miRNA, protein | 425 | 3.76E-16 | 1.68E-11 |
| Breast[4] | mRNA, miRNA, protein | 634 | 1.47E-07 | 3.56E-07 |
| Kidney[6] | mRNA, miRNA, protein | 240 | 0.91 | 0.86 |
| PCPG[5] | mRNA, miRNA, protein | 80 | 0.043 | 0.35 |
| Skin[2] | mRNA, miRNA, protein | 338 | 0.0014 | 0.0058 |
| Thyroid[1] | mRNA, miRNA, protein | 219 | 0.049 | 0.054 |
| | | | 30.21 | 23.73 |

**Supplementary table III. Comparing Spectrum's eigengap procedure with the multimodality gap on multi-omic TCGA data.** Values correspond to p values from a cox proportional hazards regression model using a log-rank test to test the significance of the survival time differences between clusters. The final row is the summed -log10(p values) for that column. PCPG stands for Pheochromocytoma and Paraganglioma.

| Dataset | Data | Samples | Eigengap | Multigap |
|---|---|---|---|---|
| Bladder[7] | mRNA, miRNA, protein | 338 | 0.0042 | 0.0028 |
| Brain[3] | mRNA, miRNA, protein | 425 | 3.76E-16 | 2.39E-16 |
| Breast[4] | mRNA, miRNA, protein | 634 | 1.47E-07 | 1.99E-07 |
| Kidney[6] | mRNA, miRNA, protein | 240 | 0.91 | 0.0019 |
| PCPG[5] | mRNA, miRNA, protein | 80 | 0.043 | 0.7 |
| Skin[2] | mRNA, miRNA, protein | 338 | 0.0014 | 0.0021 |
| Thryoid[1] | mRNA, miRNA, protein | 219 | 0.049 | 0.063 |
| | | | 30.21 | 31.63 |
